# Microbial Influenced Corrosion: A Novel Initiator of In Vivo Spine Rod Fracture

**DOI:** 10.64898/2026.06.02.729376

**Authors:** Reed Ayers, Xiaolei Guo, Christopher J. Kleck, Evan Amnidown, Jonathan Harris, Brian Gorman, Michael Rogers, David Ou-Yang, Evalina Burger, Nolan Wessell, Laura Damioli, Cheryl Ackert-Bicknell

## Abstract

Spine rod catastrophic failure/rod fracture after implantation is estimated to occur in ∼10% of all cases worldwide with this rate remaining consistent for more than 20 years. To accommodate the spine mechanical environment, metal alloys such as austenitic stainless steels, cobalt–chromium–molybdenum alloys, and α/β type titanium alloys such are used as they have a high resistance to fatigue failure. The work presented herein addresses the gap in understanding about what happens in a laboratory environment versus what is happening in human patients that results in metal leaching into tissues as well as a shortened lifespan of a spine rod in a patient population that is not capable of breaking these spine rods from purely mechanical means. Eighty-five (N=85) patients (51 female – 62.9 ± 13.5 years old, 34 male - 62.7 ± 10.9 years old) who had spine revision surgery due to patient mechanical issues, such as loss of sagittal or coronal balance, pain, pedicle screw loosening, hardware associated infection were included in this study. All explanted rods had optical indications of surface modification that were not the result of mechanical damage, e.g. gouges, scratches, notches. The presence of Ti6Al4V rods increases the presence of local tissue concentrations of elemental Ti (190.98 ± 207.84 μg/g (ppm)) in all patients over the amount that would normally be present in patients who never had any titanium based orthopedic device implants (spinous muscle, 13.12 ± 11.43 μg/g). PQS, HHQ, and NHQ ToF-SIMS molecular signals were found co-located to corrosion pits as well with AHLs and palmitic acids indicating microbial presence and metabolism in all patients. This suggests that MIC can cause in vivo pitting corrosion resulting on rod failure.

## INTRODUCTION

The use of metal alloy spinal implants enables bone fusion while maintaining the stability of the spine for patient functionality. However, this is balanced with possible implant fatigue failure due to the inability to reduce the normal cyclic loading environment of the spine.^1^ To accommodate the spine mechanical environment, metal alloys such as austenitic stainless steels (ASTM F138), cobalt–chromium–molybdenum alloys (ASTM F1537), and α/β type titanium alloys such as titanium–6 wt% aluminum– 4 wt% vanadium alloy with extra low intestinal oxygen (ASTM F136ELI) are used as they have a high resistance to fatigue failure.^2^ They are able to withstand a large number of loading cycles (>10^6–7^) without fracturing. Along with fatigue resistance these alloys are assumed to be corrosion resistant in the human in-vivo chemical milieu and pH due to their ability to from a thin metal oxide layer that should be inert in the chemistry that exists in bone and muscle.^3^

Spine rod catastrophic failure/rod fracture after implantation is estimated to occur in ∼10% of all cases worldwide with this rate remaining consistent for more than 20 years.^4–7^ Most rod failures occur within 1-4 years of implantation meaning that the implanted rod may experience between only 10-20×10^6^ loading cycles, assuming near to normal loading activity after surgery (adjusting for age).^1^ Literature examining the fatigue properties of Ti6Al4V with various microstructures suggest that dual phase Ti6Al4V spine rods should have an infinite lifespan so long as the stress imparted does not exceed approximately 220MPa (36,000 psi).^8, 9^ This suggests that titanium alloy rods in patients are breaking long before their design expectations.^10^

Previous spine rod failure work shows that local microscopic changes to the surface topology/microstructure can result in their early failure.^11^ It is also established knowledge that titanium is highly susceptible to pitting corrosion which occurs in multiple environments.^12–15^ Pitting of the Ti6Al4V surface creates micron sized local stressors that concentrate loads applied during normal function.^16^ Mathematically, Hoepner (1979) created a model for the formation of a crack resulting from coalesence of these corrosion pits, laying the foundation of how pitting corrosion leads to cracks and subsequent catastrophic fracture.^17^ Within this model, it was shown that fatigue fracture in a pitting environment can happen over 100 times faster at stresses up to 80% less than the endurance limit specific to an alloy tested in standard conditions.^18^

Studies in petrochemical, marine, power generation, manufacturing, aviation settings, show titanium alloy pitting from microbial influenced corrosion (MIC) combined with cyclic loading (fatigue) leads to early failure.^14–16, 19, 20^ Estimates of the economic costs of repair and replacement of parts failing due to corrosion is on the order of 4.2% of the American GDP with MIC causing up to 20% of the observed corrosion in any environment.^21^ By design, Ti6Al4V should not corrode substantially in the human chemical environment.^16, 22–24^ However, as the understanding of tissue microbiomes has increased through modern genetic methods^25^, it is reasonable then to assume that in an environment where microbes are known to proliferate, MIC would be present on implanted alloys. Thus, the need to study this possibility is necessary both for patient quality of life and the national healthcare economic costs, estimated to be over $1 billion over 5 years for spine patients requiring revision.^26^ The work presented herein addresses the gap in understanding about what happens in human patients that results in metal leaching into tissues as well as a shortened spine rod lifespan in a patient population that is not capable of breaking spine rods by purely mechanical means.

## RESULTS

### Patient Demographic

Eighty-five (N=85) patients (51 female – 62.9 ± 13.5 years old, 34 male - 62.7 ± 10.9 years old) who had spine revision surgery due to patient mechanical issues, such as loss of sagittal or coronal balance, pain, pedicle screw loosening, hardware associated infection were included in this study. Inclusion criteria required sufficient discarded tissue adjacent to the rods and pedicle screws for chemical analysis; hardware was in lumbar spine but could bridge into the thoracic spine; all hardware, including rods, screws, connectors were removed. Cervical hardware was excluded. All patients had tissue adjacent to the rods sent to pathology for histologic analysis and clinical microbiology for culture of any pathogens as part of the standard of care. Average time of implantation is 4.69 ± 5.00 years with no significant difference between sexes. Broken rods (N=13) or broken pedicle screws (N=2) (female 6 broken rods and 1 broken screw; male 7 broken rods 1 broken screw) were part of this patient demographic as they were one of the mechanical reasons for surgical intervention. There were 19 cases of infection (9 female; 10 male) as defined by the clinical culture of bacteria either diagnosed as bacteriemia prior to surgery or from tissues taken during surgery. Of the patients with infection, four had broken instrumentation, 3 with broken rods and 1 with a broken screw. The broken instrumentation patient demographic is typical for the size of patients at this age (65.25 ± 10.04 years; 85.80 ± 15.71 kg; 31.47 ± 5.36 BMI).

### General Appearance of Explanted Spine Rods

Upon explantation, rods were cleaned and imaged using optical microscopy and scanning electron microscopy (SEM). Broken hardware had the fracture surfaces imaged under an optical microscope as well. While most rods do not appear as Figure 1A on explantation, all rods explanted had optical indications of surface modification that were not the result of mechanical damage, e.g. gouges, scratches, notches. This data is presented in Hutchings et al., 2025.^23^ Color differences of surfaces exposed to tissues were clear, meaning a chemical modification to the rods’ passivation layer had occurred changing the surface color from the original.^27^ Figure 1B shows a representative highly corroded rod encased in bone from a patient with clinically diagnosed Candida albicans infection. Being encased in bone enabled a careful cross-sectioning with minimal chemical modification to the tissue, e.g. dehydration, preservative infiltration, to allow physical and elemental observation of the bone/alloy interface. Energy dispersive X-ray spectroscopy (EDAX) (TESCAN S8252G with EDAX Octane Elect Plus) showed how elemental titanium and vanadium were leaching from the surface of the rod into the surrounding tissue. The boney nature of the surrounding tissue is verified through the co-localized presence of phosphorous and calcium. In this specific case, the concentration of titanium and vanadium in the tissue was 163.0 μg/g and 0.8 μg/g respectively. Candida albicans has been shown to induce titanium pitting corrosion.^28^

**Figure 1:**
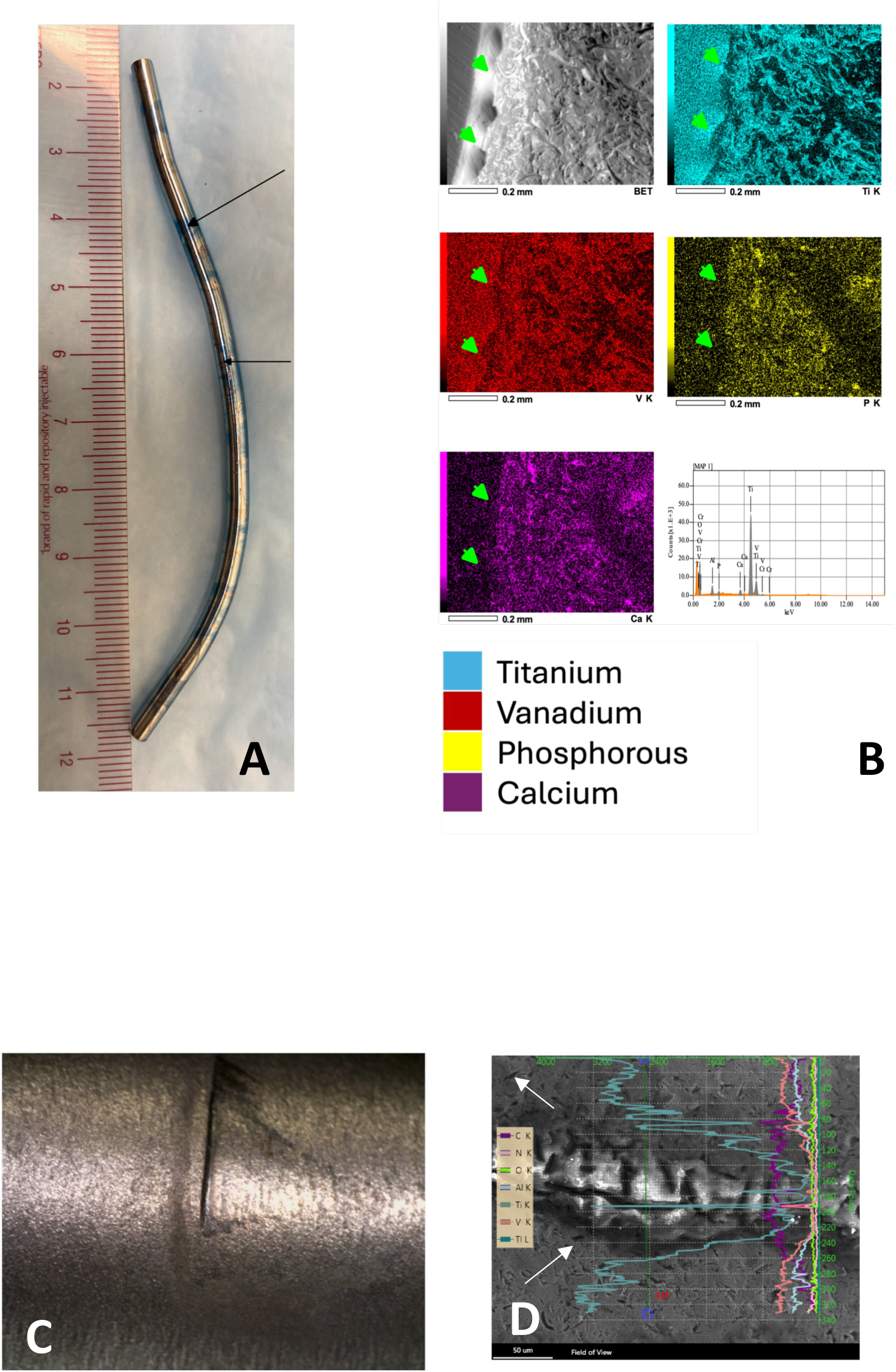
A) Image taken in the operating room of a retrieved spine rod showing extensive surface modification. The original color was a deep blue (black arrows). No evidence of rod loosening is present. Evidence of a loose rod would present as gouges, notches, and scratches on the rod surface. None were evident. B) EDS elemental mapping of a cross section of a spine rod (green arrowheads in each panel) that was encased in bone from a patient with a Candida albicans fungal infection. Leaching of Ti and V elements into the surrounding bone are documented. The individual colors correspond to Ti, V, P, and Ca respectively with the EDS images shown. C) A nascent crack on a Ti6Al4V alloy spine rod (approx. 180 micron depth) with local evidence of corrosion. D) The same crack imaged using SEM-EDAX in a backscatter mode shows the increased presence of carbon surrounding the crack with a decrease in Ti, Al, V signals. Pitting corrosion of the alloy can also be seen surrounding the crack (white arrows).

In patients with broken rods, if there is a bilateral intact rod it is also replaced as part of the standard of care. In a patient included here the partner rod to a fractured rod had a nascent crack with localized corrosion which was discovered during the physical examination, Figure 1C. This crack was approximately 180 microns in depth based on measurements made on a Zeiss Versa 520 XCT. Using a SEM in backscatter mode with EDAX mapping of the rod surface around the crack showed elemental titanium signal being obscured by organic carbon at its edges while the titanium signal is clear within the crack and outside of the biofilm present. Surface corrosion can be clearly seen from the discoloration of the titanium, and when observed under the SEM pitting is clearly present (Figure 1D).

### Quantifying Metal Leaching into Tissues Using ICP-MS/AES

Discarded tissues adjacent to the hardware were analyzed at a commercial laboratory, Hazen Analytic Laboratory, Golden, CO, USA, using inductively coupled plasma mass spectrometry (ICP-MS) (Figure 2). The presence of Ti6Al4V rods increases the presence of local spinous muscle tissue concentrations of elemental Ti (190.98 ± 207.84 μg/g (ppm)) in all patients is over the amount that would normally be present in patients who never had any titanium based orthopedic device implants (spinous muscle, 13.12 ± 11.43 μg/g). At the same time, there is a significant difference between elemental titanium concentrations versus elemental vanadium (33.94 ± 56.06 μg/g) concentrations (p=0.004) suggesting that what is present in the tissue is not particulate but rather leaching ions. The industry allowed vanadium average concentration in Ti6Al4V is 4 wt% ± 0.5wt% per ASTM F136 for Ti6Al4VELI alloys used in medicine, a ratio of 1:22.5 V:Ti, where the measured ICP ratio of V to Ti in the tissue is 17% (1:6 V:Ti). While not studied here, there may be a selective leaching of vanadium from the implant or an unknown biologic sequestration of V in the tissues. The challenge of ICP-MS on tissue is the high variability of the data due to complexing of other elements common in tissue that obscure ^48^Ti and ^51^V spectra such as ^48^Ca interfering with ^48^Ti, and ^35^Cl^16^O^+^ or ^34^S^16^OH^+^ interfering with ^51^V in biologic samples.^29^

**Figure 2:**
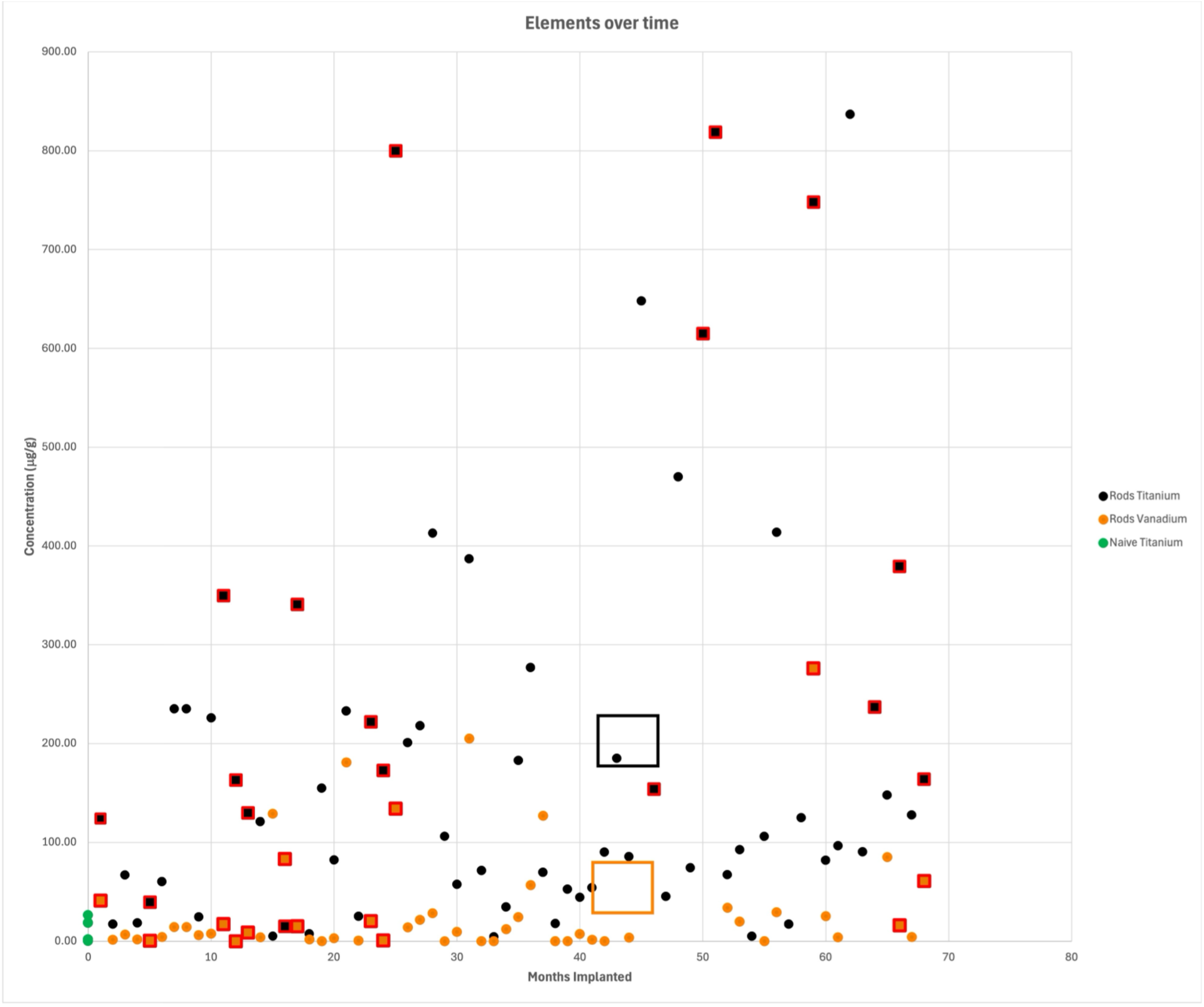
ICP-MS/AES of discarded tissues adjacent to the spine rods. The presence of rods significantly increases the concentration of tissue titanium over the value from patients with no metal implants(green dots) (p=0.004). Vanadium concentration does not mimic the titanium values, suggesting the measured concentrations are not particulates caused by wear.

The trends of an increased Ti and V presence in patients diagnosed with infection provides a possible hint as to other mechanisms resulting in implant degradation. Infection diagnosed patients had 270.66 ± 286.84 μg/g Ti tissue concentration compared to non-infected patients with 152.84 ± 167.79 μg/g suggesting that the presence of microbes in the tissue adjacent to the implant may be reflective of conditions on the rod surface.

### High Resolution Chemical Analysis of Implant Surfaces Using Time of Flight – Secondary Ion Mass Spectroscopy (ToF-SIMS)

High resolution, direct chemical observation of microbial presence in areas of surface corrosion was accomplished using Time of Flight-Secondary Ion Mass Spectroscopy (ToF-SIMS) on rods from 14 patients (10 without clinical diagnosed infection, 4 with clinical diagnosed infection). The explanted rods were sonicated to remove most blood and unbound biomaterials. Before placement in the ToF-SIMS, all rods were rinsed with ultra-pure water and ammonium formate to remove salts known to suppress TOF-SIMS signals; and due to the roughness of the sample surface, exposure to atmospheric conditions, and handling, all scanned areas were sputter cleaned using an Argon Gas Cluster source to remove any species that may not be native to the surface, such as, adventitious carbon, hydrocarbons, aldehydes, amines, and oxidation. These steps ensured that any molecules present were chemisorbed to the rod surface and unlikely to be environmental contamination.

Molecules unique to bacterial function and structure were targeted for imaging, specifically molecules associated with microbial communication that are not likely to be present in a human (Figure 3a and 3b). Both quinolones and N-acyl-homoserine lactones (AHLs), also referred to as autoinducers, enable microbial communication through quorum sensing both for bacteria and fungi.^30, 31^ Quinolones and their precursors (2-hptyl-3-hydroxyquinolone – PQS; 2- heptyl-4-quinolone – HHQ; 2-nonyl-4-hydroxyquinoline (NHQ)) and AHLs were selected because of their expressly unique microbial function.^32–34^ Quinolones are produced by bacteria in the orders of Pseudomonas, Burkholderia, as well as Shewanella and are not synthesized in humans.^35, 36^ AHL is a class of molecules separate and distinct from the quinolone quorum sensing molecules and utilized by other in other phyla such as Firmicutes for communication. AHL molecules have been well studied in previous in vitro work thus, their presence can be an indicator of microbial activity.^35, 37, 38^ Palmitic acid was selected as the third microbial molecule for analysis as it is a major structural fatty acid in microbial membranes.^39^ Palmitic acid, however, is present, but not synthesized normally in the human body and is also a component of human cellular structures. However, in this work there was little evidence of this molecule outside of corrosion regions nor in a significant amount that was not also associated with quinolones and AHLs. Baseline spectra for all molecules were taken from the literature.^33, 35, 37, 38, 40^ In this work, the presence of all three in a chemisorbed in a specific location was considered an indicator of the presence of local microbial metabolism.

**Figure 3:**
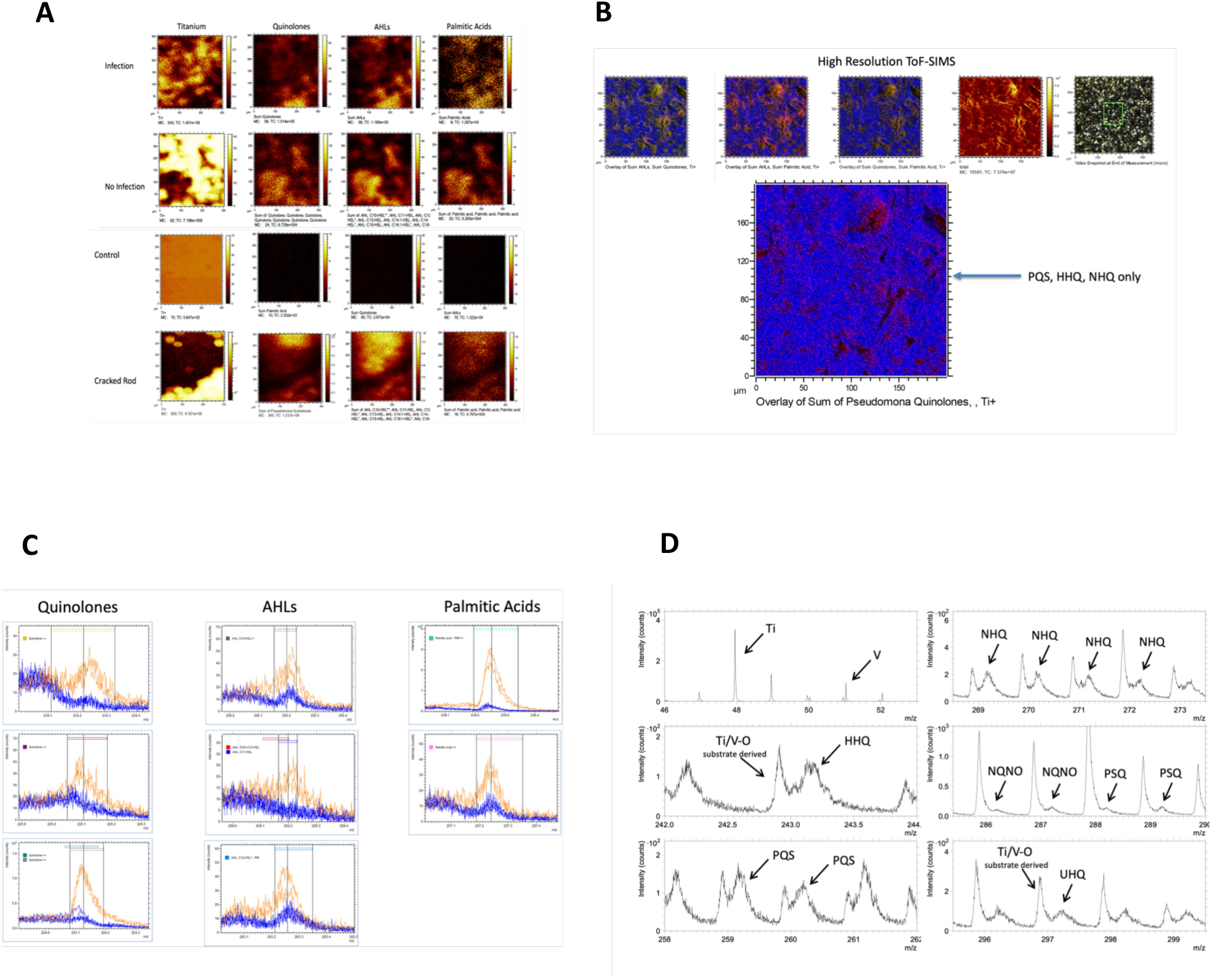
TOF-SIMS of observable corrosion areas. There is bacterial presence regardless of patient clinical infection status where there is corrosion. The colocation of autoinducer molecules and structural fatty acids is evident in regions where the Ti signal is diminished. The organic molecule images are the summation of all spectral peaks for any quinolone, AHL, or palmitic acid. A) Individual spectral images for the specific molecules. Regardless of patient infection status, the presence of colocated molecules in regions of corrosion are evident. No moecules of interest are evident on the control rods. B) High resolution ToF-SIMS shows the AHLs, quinolones, and palmitic acid molecules colocated in areas of pitting in the corrosion regions. C) Representative spectra showing comparison of corrosions (Orange) regions and adjacent uncorroded regions (Blue). The signals for the molecules of interest are not present in uncorroded regions. D) ToF-SIMS quinolone spectra showing the differences between the titania and vanadia masses are separated from the PQS, NHQ, HHQ, NQNO molecular fragments. This provides strong evidence of the resolution of ToF-SIMS to detect biologic fragments without interference from metal oxides in the same mass region.

The aforementioned molecules were present in conjunction with pitting corrosion on rods from both infected and non-infected patients (Figure 3A). In those areas, verified by SEM, PQS, HHQ, and NHQ signals are found co-located to corrosion pits as well with AHLs and palmitic acids. Depth of quinolone chemisorption into the polyoxide surface was approximately 3 nm as measured by argon beam sputtering of the surface between scans. The colocalization of all three molecules around corrosion pits suggests a non-random presence of microbes in areas of corrosion while the depth of quinolones presence suggests a chemical interaction with the surface polyoxide. The absolute number of detections of these molecules varies greatly between the corrosion areas and reference areas (Figure 3B) as well. Even when we detected quinolone, AHL, and palmitic acid in reference areas, all three molecules were never co-located. None of these signals were detected on exemplar rods (never implanted). (Figure 3A)

Because of the uniqueness of quinolones and their relationship explicitly to bacteria the identification of specific quinolone peaks is shown in Figure 3C. Labeled broad peaks correspond to quinolone molecular ions. These signals exhibit distinct spatial localization and adduct patterns (272.200, 294.182, 310.156 m/z) that are not observed in control Ti–V surfaces. In contrast, the tall, narrow peaks arise from substrate-derived Ti/V–O cluster ions, which are intrinsic to the metallic surface and remain constant across all samples. The broad peak morphology is consistent with organic molecular ions analyzed in delayed- extraction imaging mode, in which secondary-ion energy spreads naturally produce slightly wider line shapes. The consistent spatial and spectral separation between the two signal families therefore rules out post-source decay or instrumental broadening as causes for the broad NHQ peaks.

The rarity of microbial metabolic products in reference areas is evident in Figure 3D showing the representative peak counts of quinolone, AHL, and palmitic molecules in corrosion regions relative to uncorroded regions. Their presence is distinct even while the counts remain low compared to other elements and molecules. This suggests that the local corrosion is likely a function of a small number of microbes not clinically cultures and not that of pathogens cultured in clinical infections.

Principle component analysis (PCA) of the spectra yields intriguing results. Because of the untargeted nature of this ToF-SIMS analysis, a Z-score normalized spectra comparison was initially applied from corrosion and the adjacent uncorroded areas on in the retrieved rods (Figure 4). A clear difference between the spectra obtained between corrosion reigions and uncorroded regions is apparent. Peaks associated with PC1 (Table 1) and PC2 were suppressed in areas of no corrosion. Succinctly, PC1 is predominately quinolone fragments while PC2 is predominately AHL fragments with PC3 being composed of alloy surface peaks. PC1 and PC2 account for 72% of the variance in measured spectra (Figure 3E). None of the fragments present in PC1 and PC2 were detected on exemplar (never implanted) rods that were hospital sterilized prior to ToF-SIMS.

**Figure 4:**
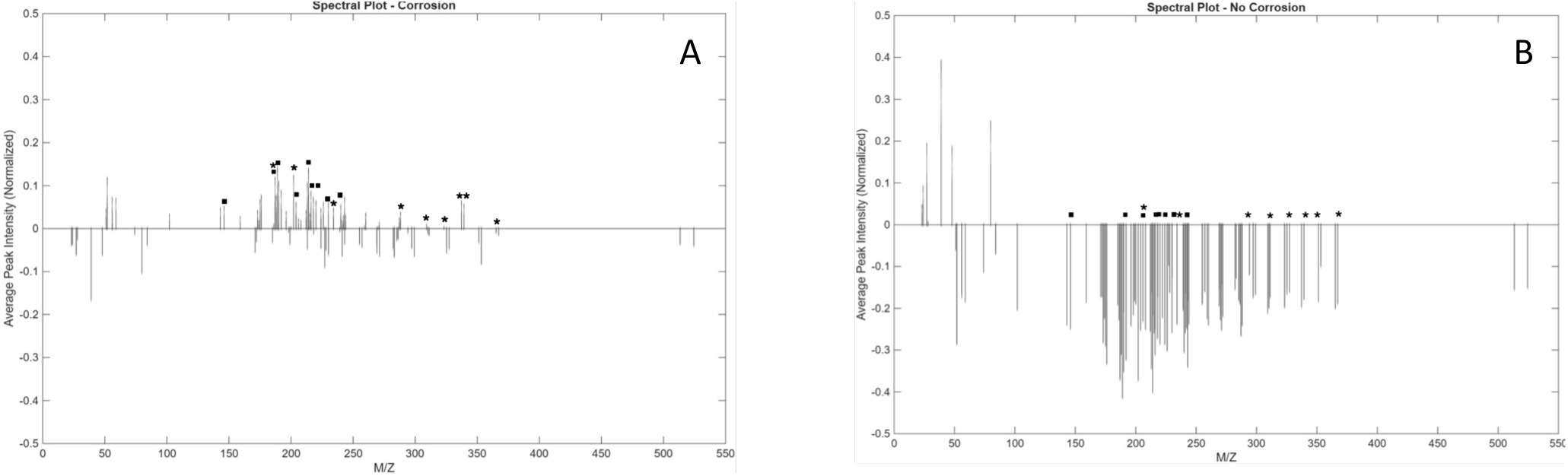
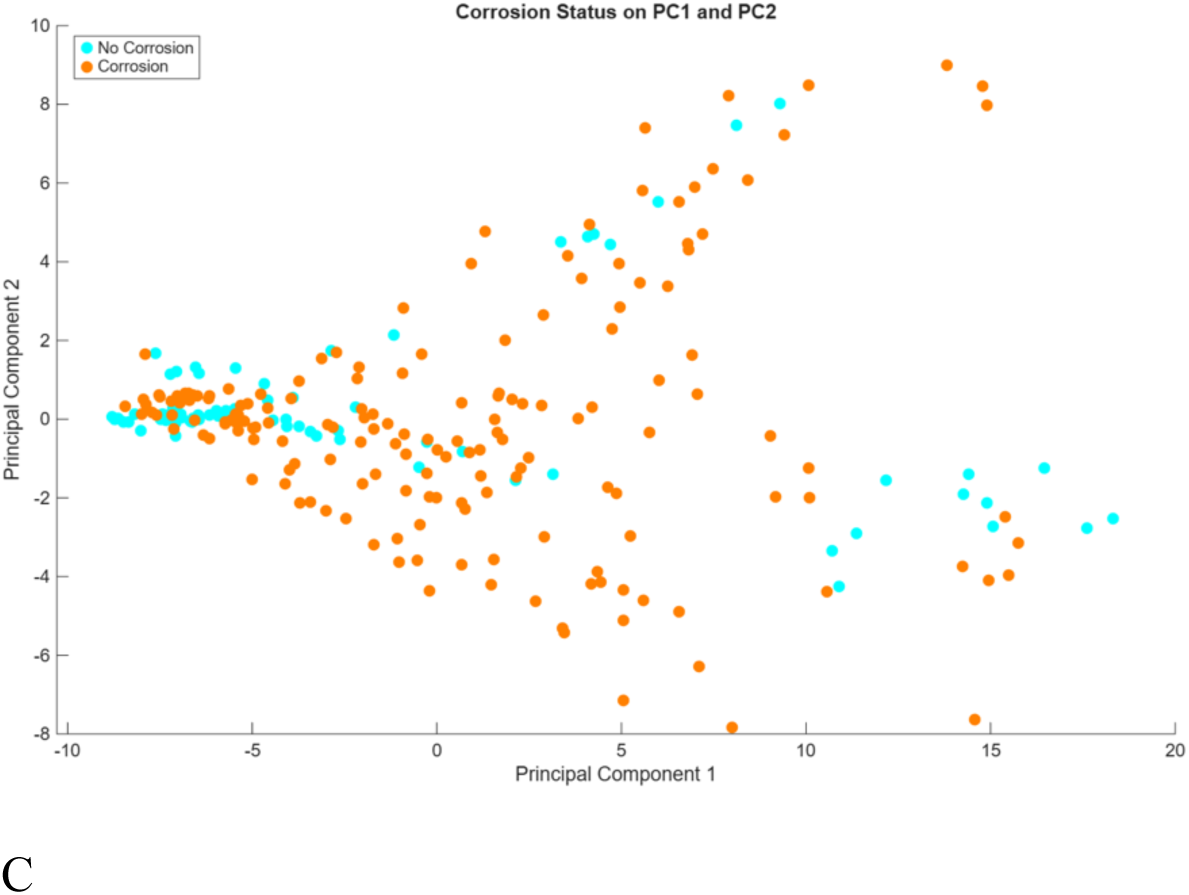
Z-score normalized spectra for the ToF-SIMS data in regions of corrosion (A) and adjacent uncorroded regions (B). ▪ Denotes fragment peaks that constitute PC1* Denotes fragment peaks that constitute PC2. 72% of the variance between the spectra are described by PC1 and PC2.

**Table 1.** Peaks Describing the First Three ToF-SIMS Spectra Principal Components.

| Principal Component 1 (PC1)<br>Fragment (m/z) | Principal Component 2 (PC2)<br>(m/z) | Principal Component 3 (PC3)<br>(m/z) |
| --- | --- | --- |
| Quinolone (218.0552) | AHL: C16-HSL (339.2587) | Mg+(23.9851) |
| Quinolone (230.0419) | AHL: C16:1-HSL (337.2202) | AHL Frag (74.0623) |
| AHL: C2-HSL (143.0615) | Quinolone+Na (NHQ)<br>(294.1796)' | AHL Frag (84.0443) |
| Quinolone (216.0408) | AHL: C14-HSL- PMI(311.2308) | AHL: C7-HSL (213.0552) |
| Quinolone (216.1346) | Quinolone (202.0484) | Al (26.9821) |
| Quinolone (196.0761) | AHL: C14:1-HSL ++ (309.2136) | Quinolone - PMI (189.0473) |
| Quinolone (188.059) | AHL: OXO-C14:1-HSL (323.2519) | Ti (47.9504) |
| Quinolone (192.0574) | AHL: C18:1-HSL (365.231) | AHL: OH-C4-HSL (187.0923) |
| Quinolone H (HHQ)<br>(244.1056) | Quinolone - PMI ++ (189.0473) | Quinolone (202.0484) |
| Quinolone (204.048) | Palmitic acid - PMI (239.1948) | Co (58.9571) |

## Discussion

Corrosion of orthopedic devices in the human has been studied predominately as an observational exercise based on fundamental corrosion concepts established in materials failure science.^41, 42^ There remains a dearth of studies that directly examine titanium alloy corrosion in vivo to characterize and understand how the in vivo environment affects the implanted alloys.^43^ The relevance of this is that the local human immune and microbiome environment is so unique as to not be able to be duplicated. This leaves a gap in knowledge in how spine rods can still fracture in use even with improvements in alloy processing and design. This work introduces a new mechanism for early spine rod failure that fits with current MIC work and explains how spine rods can break in an already mechanically compromised patient population.

Our ToF-SIMS data indicates quinolones, AHLs, and palmitic acids are collocated with corrosion pits in all patients. Quinolones themselves are not synthesized in the human and consequently they should not be present near corrosion pits (Figure 3D). The implication of this is that there is a colonization of titanium implant surfaces as part of a normal biologic integration and that the local microbial biofilm the is modifying the surface. A possibility is that these molecules are associated not just with quorum sensing but electron exchange between the microbes and the surrounding environment.^44–46^ PQS has been demonstrated to complex with vanadate (VO4^3−^) and free aluminum (Al^3+^) providing possible, although unstudied mechanisms by which corrosion occurs in vivo.^46^ In addition, the pseudomonas siderophore, pyoverdine, will form covalent bonds with TiO_2_ and AlOOH which are also present on the implanted rod surfaces^47^ Thus, we must accept that MIC spine rod pitting corrosion is ubiquitous across patients and is not a function of infection but there is likely a microbial function occurring.

The correlation of quinolone and AHL fragments to corrosion and uncorroded regions is of specific interest. While the presence of the fragments is reduced in uncorroded regions, Figure 3E suggests that their presence may be related to the microbes present. As noted prior, quinolones are produced by specific microbes and are not ubiquitous. The microbes that produce these molecules may be more capable of inducing local corrosion while microbes that communicate using AHLs may provide a protective function as indicated by the stronger association of PC2 with uncorroded regions.

To discover sources of quinolone and AHL metabolites attempts were made to characterize the microbes present in these areas. Hardware was sonicated to remove the biofilms for bacterial DNA identification (N=30). However the extremely low DNA mass prevented consistent extraction and significant measurement on a per patient basis (N=4). While patients yielded sequencable bacterial DNA, in 26 cases the sequencing reads were below accepted limits of statistical significance. The samples that produced sufficient DNA mass to sequence, possible sources of quinolones arose as well as showing a diverse microbial presence on hardware (Figure 5). These patients showed the presence of Burkholderiales order bacteria (in the Betaproteobacteria class) with the family of Alcaligenaceae. This is significant as the Alcaligenaceae family bacteria produce quinolones as part of their metabolism similar to other microbes in the Pseudomonadota phylum.^48, 49^ As previously described, the quinolones produced can utilize the surface metal oxides providing a source of multivalent metal ions for bacterial metabolism as well as degrading the local surface, resulting in corrosion pits. While not a significant measure for this study, this data suggests that these microbes are present on all spine rods regardless of patient infection status which further bolsters the hypothesis that MIC is occurring in vivo. Subsequent medical and biologic studies will need to

**Figure 5.**
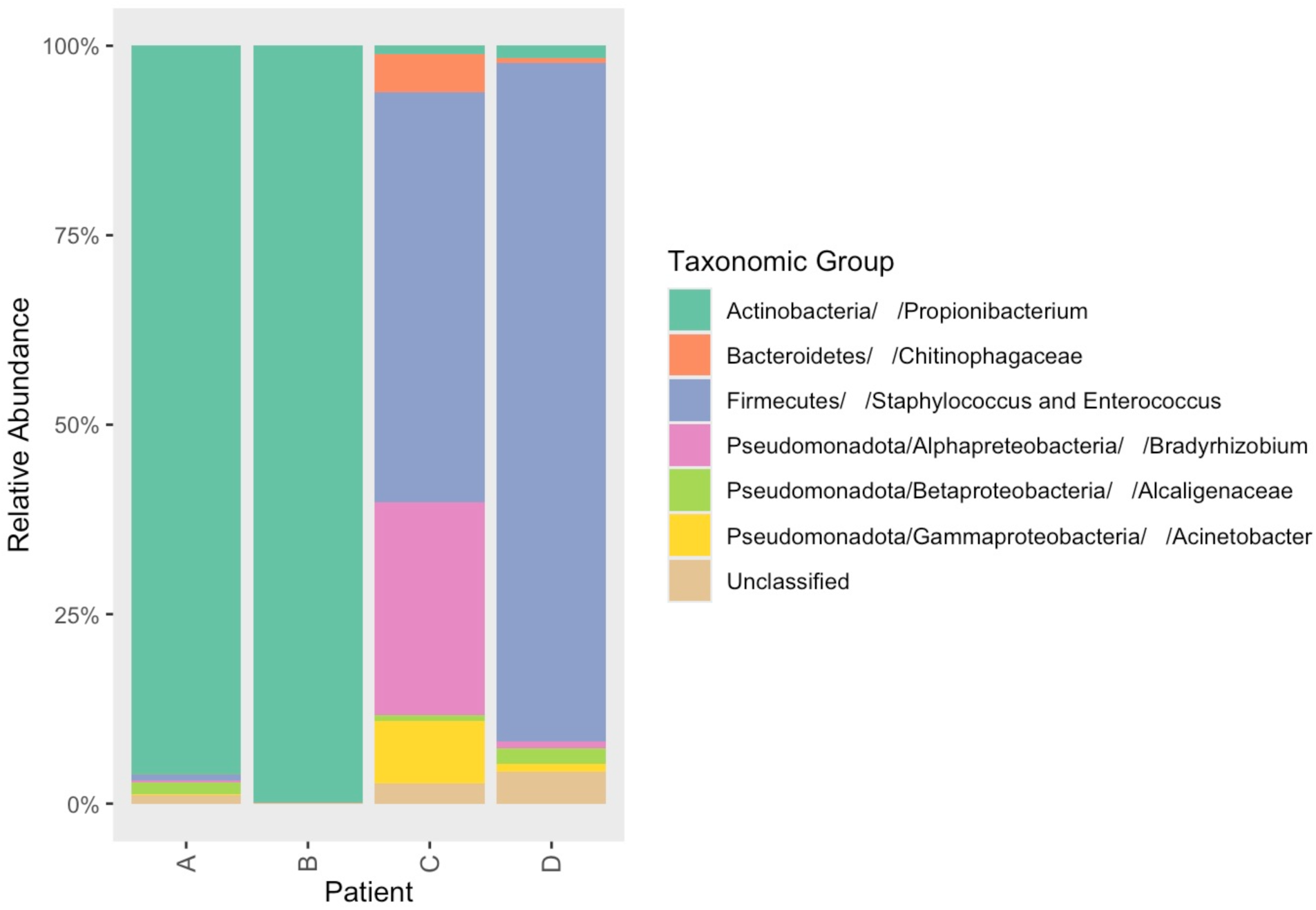
Pateint A Infection (broken Rod) C. acnes clinical diag. Patient B no infection C. acnes primary microbe detected with Staphylococcus/Enterococcus and Alcaligenaceae also present Patient C Polymicrobial Infection (Broken Rod) Pateint D Infection Staph a. clinical diag

## Methods

The retrieval of the spine rods used in this work was accomplished under an institutionally approved research protocol. Patients were identified prior to revision spine surgery through radiographic and medical record review. Patient inclusion criteria was the presence of instrumented thoracic/lumbar regions regardless of diagnosis and age 18-85 years old. Exclusion criteria was patient pregnancy, cancer or other diseases where immunosuppressive agents are utilized. Consent for this research was obtained prior to patient revision procedure.

Upon explantation all spine hardware was placed in sterile containers with sterile Dulbeccos phosphate buffered solution (PBS) and sealed for sonication. Pedicle screws and associated hardware were stored and sonicated separately from rods. Sonication was conducted in a sterile biosafety hood using a Branson sonicator (Branson Model 3800 CPX) at 40kHz for 25 minutes. Sonicate fluid was decanted into 50ml centrifuge tubes, vortexed for 5 seconds and then centrifuged for 10 minutes at 1,000g. All but 2 ml of sonicate fluid is decanted and disposed with the remaining frozen at -20°C in preparation for 16s metataxanomic analysis.

Observation of corroded and uncorroded regions was conducted using SEM (TESCAN S8252G, Libušina Kohoutovice, Czech Republic) with local elemental constituency quantified by EDAX (Octane Elect Plus, EDAX, LLC, Pleasanton, CA, USA) in regions of interest, based upon presence of macroscopically observable surface damage. Areas of discoloration as well as what appeared to be uncorroded regions were imaged. Energy dispersive spectroscopy (EDAX) was utilized to verify rods were Ti6Al4V alloy and if corrosion products were present in on the surface.

### ToF-SIMS

Prior to testing, samples were subjected to a surface cleaning procedure to minimize extraneous contamination. Each sample was rinsed in an aqueous ammonium formate solution to remove residual salts that could suppress the secondary ion signal of the species of interest. Following the rinse, samples were gently wiped with cleanroom-grade wipes and promptly transferred to the load-lock chamber of the TOF-SIMS instrument and placed under high vacuum to minimize atmospheric exposure prior to analysis.

Secondary ion mass spectrometry was performed using a TOF-SIMS V (IONTOF GmbH, Münster, Germany) equipped with a 30keV bismuth liquid metal ion gun (LMIG). Data were acquired with a Bi₃⁺ primary ion source operated at ∼0.4 pA at 200us cycle time and rastered across the regions of interest. Spectra were collected in both positive and negative polarity modes to capture complementary information on cationic and anionic species.

To further reduce surface contamination and improve signal from underlying layers, samples were sputter-cleaned using an argon gas cluster ion beam (GCIB) operated at 10 keV with clusters containing ∼1,200+ atoms. The sputter beam was applied at ∼8 nA and rastered across the analysis area for 2–6 cycles, depending on the surface roughness of each sample. Post- sputter analyses were performed under identical acquisition conditions to enable direct comparison with pre-sputter data.

300 µm by 300 µm areas of explanted spinal rods were scanned using ToF-SIMS. Areas were selected to be either entirely an area of corrosion or an area free from visible corrosion using the 1x zoom on the optical microscope portion of the ToF-SIMS. Multiple areas of visible pitting corrosion and areas free from visible corrosion were scanned on each rod. Because of the distance between them, the areas were assumed to be independent and therefore each scan was treated as an independent observation. 16 rods from 7 different patients were scanned. The environment of each rod was assumed to be independent of the other rods, even if the rods were explanted from the same patient. Therefore, the scans from each rod were treated as independent observations. In one case, two sets of rods came from the same patient from two different surgeries. Because of the length of time between surgeries, each scan on these rods were considered independent observations.

Data from each scanned area (with and without pitting corrosion) for all rods from all patients were compiled together for 105 peaks of interest, with m/z values ranging from ∼22 to ∼524. This dataset was then additionally separated into two additional datasets: one for all scans of areas of visible pitting corrosion, and one for all scans of areas without visible pitting corrosion. The datasets were each normalized prior to performing analysis, following common practice for performing multi-variate analysis of ToF-SIMS data.

Principal Component Analysis (PCA) was performed on each normalized dataset in MatLab R2026a using the PCA function of the Statistics and Machine Learning Toolbox. The number of components that explained 95% of the variance was calculated. K-Means Clustering was then performed using ‘sqeuclidean’ as the method with 10 replicates of each dataset. The PCA scores (the location of each observation on each principal component) were the input data for the K-Means calculations and used the number of principal components explaining 95% of the variance as the initial guess for the number of clusters. The number of clusters was then refined using a silhouette plot.

### Inductively Coupled Plasma Mass Spectroscopy (ICP-MS)

A local fee for service commercial lab with the ability to conduct ICP-MS measurements (Perkin Elmer NexION 300D located at Huffman Hazen Laboratories, Wheat Ridge, CO) was used to measure tissue metal concentrations (ppb levels) from implant adjacent tissues taken at the time of surgery. All tissue was within less than 1cm distance from the implant, placed in Teflon digestion tubes. Tissues are digested in

### 16 Microbiome profiling

DNA extraction is performed using the Qiagen EZ1 Advanced kit per manufacturer’s instructions. Total bacterial load is determined using qPCR amplification of the V1/V2 region of the 16S rRNA gene and bacterial profiles will be determined by sequence analysis of 16S rDNA. Taxonomic information is assigned using the SINA classifier based on Silva reference sequences after appropriate sequence quality checks.

### Statistical Analysis

For each sample, we will calculate the Alpha Diversity, which is a measure of the richness (number of species) detected in the sonicate using the Shannon diversity index. Because of the non-parametric nature of the Shannon index, we will test for difference in Alpha diversity between cases and contols using a repeated measure linear regression. For association of metallosis with individual taxa, we will use a χ^2^ test. Beta diversity will be calculated to capture the richness/evenness between types of samples (rods, screws). We are not able to estimate power for this aim using our data (n=2), but simulations of microbiome data conducted by others suggests that with an n=60, we will have ample power to detect a difference in Beta diversity between cases and controls. The association between bacteria present (at the level of family) and measures of corrosion from Aim 1 will be determined by Pearson’s Correlation as described previously.

Figure – Z-score Comparison of spectra Averaged from regions with corrosion and regions with no corrosion. Normalized Peak Intensity Mass Spectra of Corrosion and No Corrosion Data (Top Row - Corrosion, Bottom Row - No Corrosion, Left Column - Dataset Only, Right Column - Bootstrapped Data)

Comparing the left (dataset alone) and right (bootstrapped) columns of Figure 12, we again see that the average normalized peak intensity mass spectra of the dataset alone is nearly identical to that of the bootstrapped dataset.

Even more apparent, comparing the top row (corrosion) to the bottom row (no corrosion) shows a clear mass dependence between places of corrosion and without corrosion. The M/Z region of ∼150 to 250 appears to “flip” from positive to negative for many peaks. This indicates that the presence (or a positive peak) or lack of (negative peak) is associated with a site showing corrosion or being free from corrosion. Meaning, the presence of quinolone and AHL molecules is associated with areas of corrosion, and their lack of is associated with no corrosion present.

Mass spectra were also generated using the PCA loadings for principal components 1 and 2. The bootstrap dataset shows agreement with the original dataset. The sign (positive or negative) of the peaks indicate if the presence of a molecule fragment (M/Z) is associated with a principal component (positive sign) or the lack of is associated with a principal component (negative sign). This data is for both corrosion and no corrosion because the loadings cannot be split between two groups without running a PCA on each group, which would make the loadings non-comparable

